# Direct and context-dependent effects of light, temperature, and phytoplankton shape bacterial community composition

**DOI:** 10.1101/091967

**Authors:** Sara F. Paver, Angela D. Kent

## Abstract

Species interactions, environmental conditions, and stochastic processes work in concert to bring about changes in community structure. However, the relative importance of specific factors and how their combined influence affects community composition remain largely unclear. We conducted a multi-factorial experiment to 1) disentangle the direct and interaction-mediated effects of environmental conditions and 2) augment our understanding of how environmental context modulates species interactions. We focus on a planktonic system where interactions with phytoplankton effect changes in the composition of bacterial communities, and light and temperature conditions can influence bacteria directly as well as through their interactions with phytoplankton. Epilimnetic bacteria from two humic lakes were combined with phytoplankton assemblages from each lake (“home” or “away”) or a no-phytoplankton control and incubated for 5 days under all combinations of light (surface, ∼25% surface irradiance) and temperature (5 levels from 10°C to 25°C). Observed light effects were primarily direct while phytoplankton and temperature effects on bacterial community composition were highly interdependent. The influence of temperature on aquatic bacteria was consistently mediated by phytoplankton and most pronounced for bacteria incubated with “away” phytoplankton treatments, likely due to the availability of novel phytoplankton-derived resources. The effects of phytoplankton on bacterial community composition were generally increased at higher temperatures. Incorporating mechanisms underlying the observed interdependent effects of species interactions and environmental conditions into modeling frameworks may improve our ability to forecast ecological responses to environmental change.

## Introduction

Species interactions, environmental conditions, and stochastic processes affect the abundance, diversity, and distribution of organisms in the environment. Determining the relative importance of specific factors and how they interact to structure communities remains a major challenge in ecology (Agrawal et al. 2007, Sutherland et al. 2012). Effects of environmental conditions on a population include direct effects that can potentially be characterized by single-species observations and experiments as well as indirect effects that depend on species interactions and require more community-focused approaches to detect and predict (Gilman et al. 2010). Concurrently, the outcome (e.g., mutualism, parasitism) and strength of interspecific interactions can change depending on environmental conditions as well as community composition (He et al. 2013, Chamberlain et al. 2014). To predict how communities will respond to seasonal environmental fluctuations and long-term changes in climate, it is necessary to account for species interactions that mediate and are affected by environmental conditions.

The signal of changing environmental conditions can be muted, augmented, or relayed by interactions with other species including mutualism, competition, parasitism, and food web interactions (Gilman et al. 2010). Interaction-mediated effects of environmental change have consequences for local abundances and geographic distribution as well as phenology (Miller-Rushing et al. 2010, Wisz et al. 2012, HilleRisLambers et al. 2013). For example, competitive interactions, predators, consumers, disease agents, or the absence of a mutualist partner may counteract environmental changes expected to expand the geographic range or increase the local abundance of a species (Gilman et al. 2010, HilleRisLambers et al. 2013). Additionally, shifts in the phenology of one species can result in temporal mismatch and population declines in a mutualist partner or consumer species (Winder and Schindler 2004, Edwards and Richardson 2004). In many cases, effects of environmental changes mediated by species interactions are expected to be more consequential than direct effects of change (Davis et al. 1998).

In addition to species interactions modulating the signal of environmental change, environmental conditions can alter the outcome and strength of context-dependent species interactions. A change in outcome occurs when the effect of an interaction on a given species (positive, negative, or neutral) depends on the environment. For example, the net costs of plants associating with mycorrhizae can exceed the net benefits when nutrient availability is high or light availability is low (Johnson et al. 1997). Context-dependent outcomes have been observed for competition, mutualism, and predation, but are most frequently detected for mutualisms (Chamberlain et al. 2014). In contrast, variation in interaction strength, or the magnitude of effect sizes, across contexts is relatively consistent among interaction types (Chamberlain et al. 2014). Population-level interactions can cause shifts in community structure, making it important to investigate the consequences of context-dependence at the community level.

Planktonic communities from temperate lakes are well suited for investigating interactions between biotic and environmental factors. These communities undergo annually repeated seasonal succession driven both by species interactions and environmental conditions, and respond rapidly to experimental treatments (Kent et al. 2006, 2007, Sommer et al. 2012, Weisse et al. 2016). Moreover, lakes have been described as sentinels of climate change, in part, because they respond rapidly to changes in the environment (Adrian et al. 2009). Light availability and temperature are two ecologically important factors that respond to environmental change in lakes. Increases in dissolved organic carbon concentration, as have been observed in lakes across parts of North America and Europe (Monteith et al. 2007), decrease light availability due to more rapid light attenuation and shift the vertical distribution of heat towards the surface (Bukaveckas and Robbins Forbes 2000, Read and Rose 2013). Global mean lake surface temperature has been increasing 0.34°C per decade in response to climate forcing (O’Reilly et al. 2015). Long-term climate changes are overlaid on a dynamic system where light availability and temperature change with depth and over the course of a year, especially in lakes that stratify in the summer and mix in the fall and spring.

Light and temperature can influence bacterial community composition directly as well as through bacterial interactions with phytoplankton. Wavelength-specific light attenuation in the water column creates a spectrum of niches for phototrophs (Stomp et al. 2007), including photosynthetic organisms as well as photoheterotrophs (Martínez-García et al. 2011, Evans et al. 2015). Additionally, bacterial growth is affected in a strain-specific manner by light in the ultraviolet range (Agogue et al. 2005, Hörtnagl et al. 2010). Bacteria have diverse optimal growth temperatures and ranges, such that temperature can determine outcomes of competition between bacterial populations (Upton et al. 1990, Hall et al. 2008). Phytoplankton interact with bacterioplankton through mechanisms that include selective grazing by mixotrophic phytoplankton (Flynn et al. 2012), serving as a habitat for bacterial epiphytes (Jasti et al. 2005), and providing species-specific resources as detritus (Van Hannen et al. 1999) or labile exudates released by living cells (Teeling et al. 2012, Sarmento and Gasol 2012). As temperature increases, mixotrophic phytoplankton are theorized to become more heterotrophic; this has been experimentally demonstrated with the chrysophyte *Ochromonas* sp. (Wilken et al. 2012). Additionally, the concentration and composition of extracellular organic carbon excreted by phytoplankton depend on light and temperature conditions (Zlotnik and Dubinsky 1989, Parker and Armbrust 2005).

Our objective was to characterize the combined effects of phytoplankton, light, and temperature on bacterial community composition from two humic lakes in Northern Wisconsin where interactions with phytoplankton are partially responsible for orchestrating changes in bacterial composition through time (Kent et al. 2007, Paver et al. 2013). Change in light and temperature with depth is especially pronounced in darkly stained humic lakes (Huovinen et al. 2003). We specifically aimed to determine 1) whether the influence of light and temperature on bacterial communities is mediated by phytoplankton and 2) whether the effects of interactions with phytoplankton depend on light and temperature conditions (Fig. S1). If light and temperature affect bacteria through interactions with phytoplankton, then it is expected that the variation in bacterial community composition explained by light and temperature treatment will be greater in microcosms where phytoplankton are present relative to those where phytoplankton are absent. If phytoplankton effects depend on the light and temperature context, the variation in bacterial community composition explained by phytoplankton will change under different light and temperature conditions.

## Materials and Methods

### Study sites

South Sparkling Bog (SSB; 46°00’13.6"N, 89°42’19.9"W) and Trout Bog (TB; 46°02’27.5"N, 89°41’09.6"W) are two north temperate humic lakes in Vilas County, Wisconsin that have been studied as part of the North Temperate Lakes Microbial Observatory. SSB and TB were selected for their similarity in maximum depth (∼8m) and differences in phytoplankton community composition (Paver et al. 2013). These lakes are characterized by acidic pH and high levels of dissolved organic carbon (Paver et al. 2013).

### Experimental design

We conducted a multi-factorial microcosm experiment to determine the direct and interactive effects of phytoplankton presence and composition, temperature, and light on bacterial community composition. On 6 July 2011, microorganisms were collected from SSB and TB integrated epilimnion samples (0-1m). Filtration through a 1μm Polycap AS cartridge filter (Whatman, Piscataway, NJ, USA) was used to separate bacteria from larger organisms. Phytoplankton assemblages were collected by filtering lake water through a 100μm nylon mesh (Spectrum Laboratories, Rancho Dominguez, CA, USA) to remove zooplankton and collecting, then rinsing phytoplankton cells captured on a 20μm nylon mesh with SSB water filter-sterilized through a 0.2μm Polycap AS cartridge filter (Spectrum Laboratories), which allowed smaller organisms such as heterotrophic nanoflagellates and bacteria to pass through. Phytoplankton collected on 20μm mesh were resuspended in 0.2μm filter-sterilized water from SSB, concentrating phytoplankton from 40L of lake water to 2.5L of sterilized water. All combinations of bacteria from each lake (5L of 1μm filtered water) were combined with 0.25L of concentrated phytoplankton from one of the two lakes lake or a no-phytoplankton control (0.25L of 0.2μm filter-sterilized SSB water) in triplicate 10L LDPE cubitainers (I-Chem, Rockwood, TN, USA). Combined bacteria and phytoplankton were gently inverted to mix and then partitioned into 500ml clear glass bottles (Wheaton, Millville, NJ, USA) in a predetermined, randomized order (33 bottles/ treatment).

For each bacteria-phytoplankton combination, three bottles were used to characterize the initial community composition and three bottles were incubated for five days under each of five temperatures and two light levels (Fig. 1). Temperature treatments were established and maintained by continuously pumping defined proportions of high temperature (∼25°C) surface water and low temperature (∼5°C) subsurface water (1:0, 3:1, 1:1, 1:3, 0:1) into floating plastic container incubators (0.73m × 0.53m × 0.46m, The Container Store, Coppell, TX, USA) (Fig. S2). High light and low light treatments were established by incubating bottles at the surface and bottom (∼25% of surface irradiance) of each floating container incubator. Light and temperature conditions were monitored throughout the incubation using HOBO light and temperature pendant data loggers (Onset, Pocasset, MA, USA) with three loggers placed in each container: two at opposite corners at the surface and one on the bottom.

**Figure 1.**
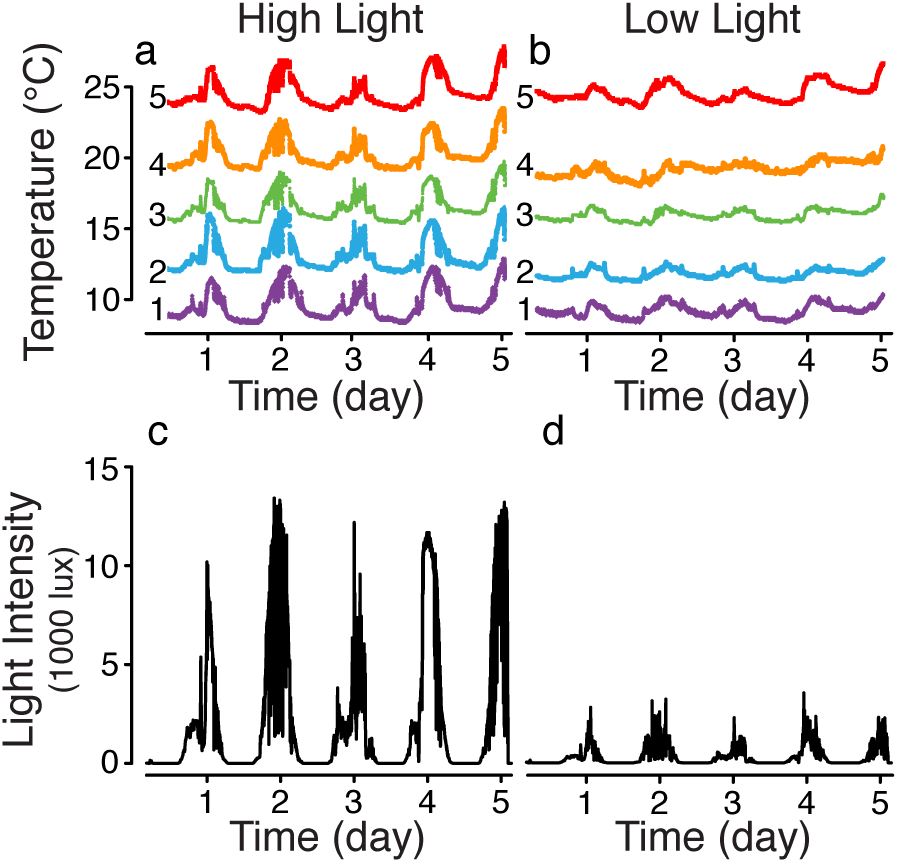
Temperature (a, b) and average light intensity (c, d) at the top (high light; a, c) and bottom (low light; b, d) of each temperature incubator over the five-day incubation. Temperature treatments are labeled 1 (coldest) to 5 (warmest).

### Microbial community analysis

Microorganisms from initial samples and from each bottle microcosm following incubation were concentrated onto 0.22μm filters (Supor-200; Pall Gelman, East Hills, NY) and frozen at −20°C. DNA was extracted using FastDNA purification kits (MP Biomedicals, Solon, OH, USA). Bacterial community composition was characterized using automated ribosomal intergenic spacer analysis (ARISA) (Fisher and Triplett 1999) as described by Paver et al. (2013). Fluorescently labeled ARISA PCR amplicons were combined with a custom 100 – 1250 bp Rhodamine X-labeled internal size standard (Bioventures, Murfreesboro, TN) and analyzed by the Keck Center for Functional Genomics at the University of Illinois via denaturing capillary electrophoresis using an ABI 3730XL Genetic Analyzer (Applied Biosystems Inc., Carlsbad, California, USA). Electropherograms from each sample were aligned and peaks greater than 500 fluorescence units were sized and grouped into bins of operational taxonomic units using GeneMarker version 1.95 (SoftGenetics, State College, PA, USA). ARISA fragments known to correspond to chloroplasts were removed from the analysis. The signal strength of each peak was normalized to account for run-to-run variations in signal detection by dividing the area of individual peaks by the total fluorescence (area) detected in each profile.

### Statistical approach

Pairwise Bray-Curtis similarities were calculated for every combination of samples using Hellinger-transformed ARISA data and visualized using non-metric multidimensional scaling in PRIMER version 6 (PRIMER-E Ltd, Plymouth Marine Laboratory, UK) (Clarke and Warwick, 2001). Permutational multivariate analysis of variance (PERMANOVA) was used to test: 1) the effects of light and temperature on bacterial community composition following incubation for each combination of bacteria and phytoplankton and 2) the effect of each phytoplankton treatment (compared to the no phytoplankton control) on bacterial community composition at each light and temperature level, stratified by the bacterial community source lake. PERMANOVA is a non-parametric multivariate analysis of variance that generates p-values using permutations (McArdle and Anderson 2001, Anderson 2001). PERMANOVA tests were run using the adonis function from the vegan package (Oksanen et al. 2011) in the R statistical environment (R Core Development Team, 2010).

## Results

### Direct and phytoplankton-mediated light and temperature effects

Over the five-day incubation, phytoplankton presence and composition, light availability, and temperature affected bacterial community composition (Fig. 2). We assessed direct effects of light and temperature on bacterial communities by analyzing changes in bacterial community composition in no-phytoplankton control treatments. Light had a significant direct effect on the composition of bacterial communities from both lakes (Fig. 3). In contrast to light, temperature had a significant direct effect on the community composition of bacteria from SSB, but not bacteria from TB (Fig. 3).

**Figure 2.**
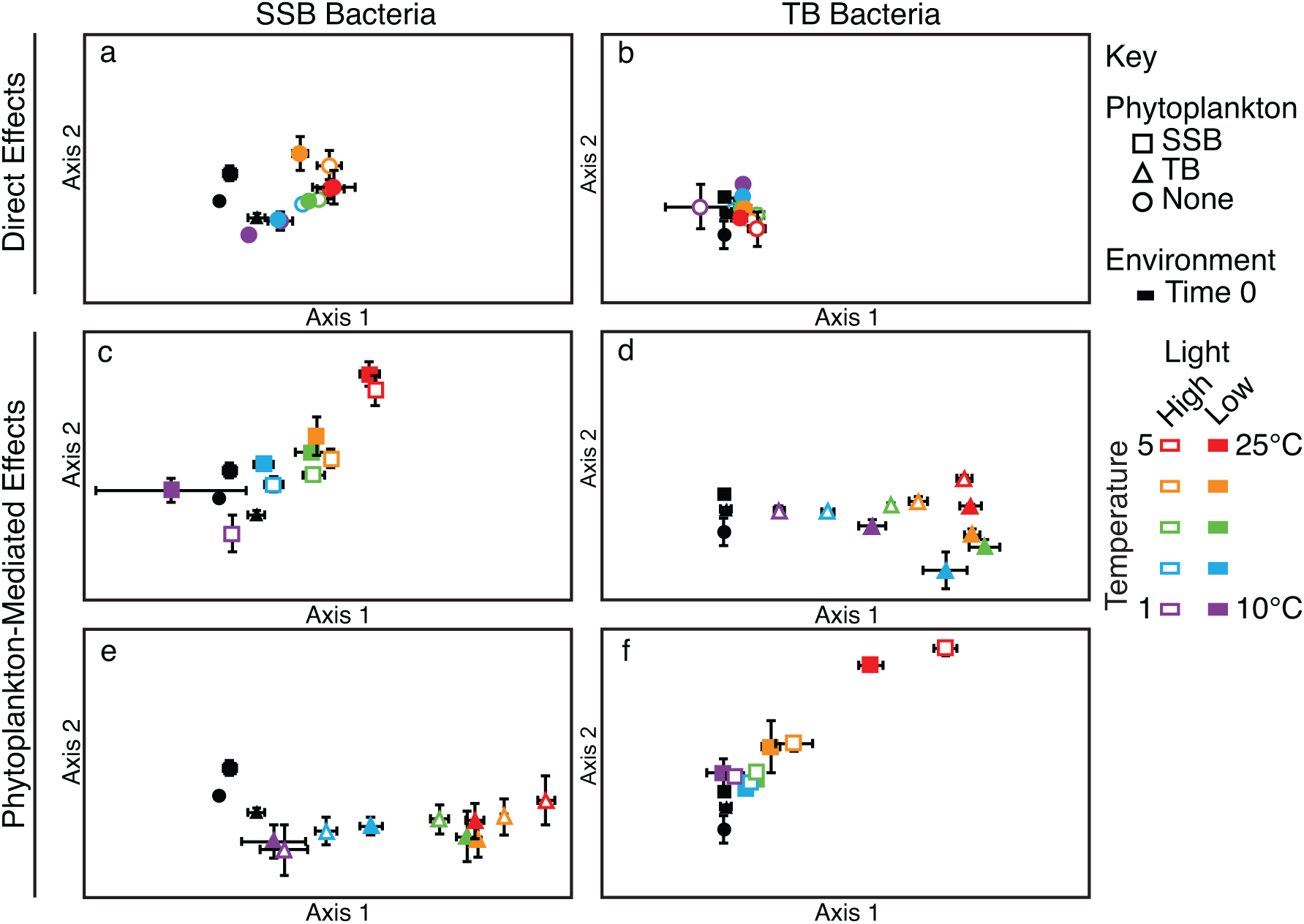
Non-metric multidimensional scaling ordination of bacterial community composition in - microcosms (average ± standard error) with SSB bacteria (a,c,e; stress value=0.08) and TB bacteria (b,d,f; stress value=0.14) before and after incubation. To simplify depiction of overlapping treatments, community composition in no-phytoplankton control microcosms following incubation is shown in plots a and b, community composition in “home” phytoplankton treatments is shown in plots c and d, and community composition in “away” phytoplankton treatments is shown in plots e and f. Community composition before incubation (Time 0) is included in all plots for reference.

**Figure 3.**
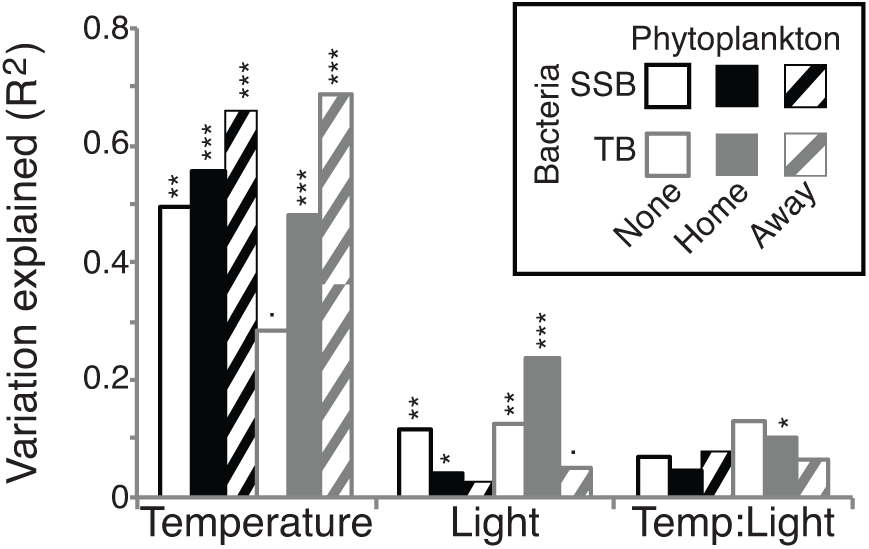
Variation in bacterial community composition explained by temperature, light, and the interaction between temperature and light for each combination of phytoplankton and bacteria. *P* values below 0.01 are indicated by symbols (*** <0.001, ** <0.01, * < 0.05,. <0.01).

The effects of light and temperature on bacteria incubated with phytoplankton depended on the specific combination of phytoplankton and bacteria (Fig. 3). When TB bacteria were incubated with their “home” phytoplankton, there was a significant interaction between light and temperature. Light explained small but significant variation in bacterial community composition when SSB bacteria were combined with their “home” phytoplankton. Significant light effects were not detected when bacteria were combined with phytoplankton from the “away” lake. In contrast, temperature had a consistently significant effect on the composition of bacterial communities when phytoplankton were present. Notably, the variation in bacterial community composition explained by temperature was higher for bacteria incubated with phytoplankton from the “away” lake than for bacteria incubated with phytoplankton from their “home” lake.

### Context-dependence of phytoplankton interactions

We used percent variation in bacterial community composition explained by phytoplankton treatment to evaluate the strength of phytoplankton effects (Fig. 4). Prior to incubation, 22% of the variation in bacterial community composition in SSB phytoplankton and corresponding control treatments was explained by phytoplankton treatment. Following incubation, variation explained by SSB phytoplankton was only greater than the initial explained variation in microcosms incubated at the highest temperature (45% variation explained). In contrast, variation in bacterial community composition in TB phytoplankton and corresponding control treatments due to phytoplankton treatment was not significant prior to incubation (p>0.05). At the coldest two temperatures, 13% and 19% more variation in bacteria community composition was explained by TB phytoplankton in low light compared to high light treatments. As temperature increased, the variation explained by TB phytoplankton generally increased until peaking at the second warmest temperature.

**Figure 4.**
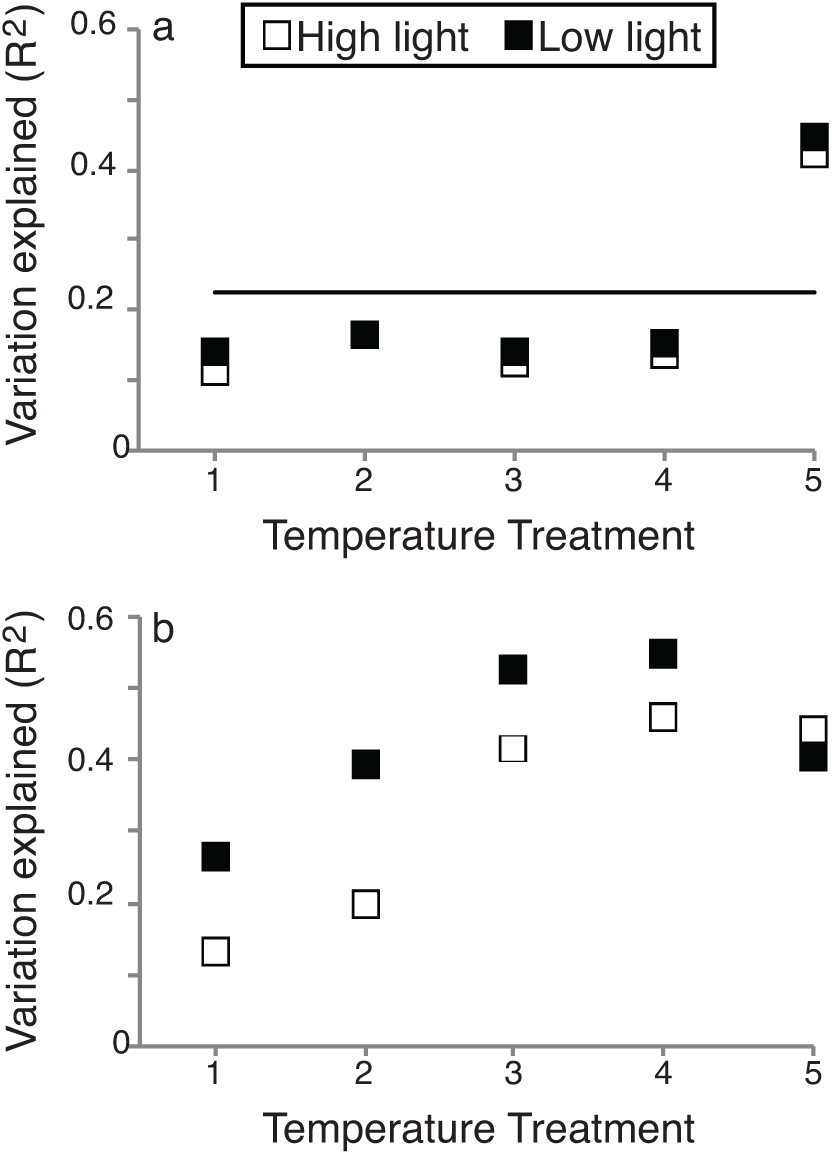
Variation in bacterial community composition explained by SSB phytoplankton treatment (a) and TB phytoplankton treatment (b) across all temperature and light conditions. A horizontal line indicates variation in bacterial community composition explained by SSB phytoplankton prior to incubation. TB phytoplankton did not explain significant variation in bacterial community composition before incubation.

## Discussion

It is well established that species interactions and environmental conditions act in concert to affect community composition, but how these factors combine to determine community composition is largely undefined. In this study we selected light, temperature, and phytoplankton-bacterial interactions to investigate the interplay among biotic and environmental factors at the community level. In general, observed treatment effects were highly similar across replicates, suggesting that deterministic processes controlled the development of bacterial communities over the course of the five-day experimental incubation. We observed direct light effects, phytoplankton-mediated temperature effects, and temperature-and light-dependent phytoplankton effects, each of which is explored below.

### Direct light effects

Light had a consistent, direct effect on bacterial community composition in experimental microcosms. One potential explanation for the direct effect of light on bacterial composition is selection for phototrophic bacteria generally, or specific types of phototrophic bacteria. Organisms that harvest light energy are adapted to use specific wavelengths of light and, as a result, phototroph distribution within the water column reflects the available light spectrum (Vila and Abella 2001, Haverkamp et al. 2008). As our study lakes have high humic content, light spectra are enriched in long wavelength photons (>600nm) and light attenuates rapidly with increased depth, creating specific spectral niches across modest changes in depth (e.g., 0.5m) (Vila et al. 1998). Photoheterotrophic bacterial cells can comprise a sizable portion of lake bacterial communities, and change in abundance with depth (Mašín et al. 2008, Martínez-García et al. 2011, Lew et al. 2016). Alternatively, or perhaps additionally, direct effects of light may have been caused by ultraviolet radiation. Based on the transparency properties of borosilicate glass, plankton were exposed to a fraction of long wave ultraviolet A radiation, but none of the ultraviolet B and shorter wavelengths of light, penetrating to their respective incubation depths (Döhring et al. 1996). Effects of ultraviolet radiation can decrease the growth efficiency of certain bacterial strains while having no effect or increasing the growth efficiency of other strains (Hörtnagl et al. 2010). It is additionally possible that products of dissolved organic matter photolysis, including low molecular weight dissolved organic matter and reactive oxygen species induced changes bacterial community composition (Glaeser et al. 2010, Paul et al. 2011).

### Phytoplankton-mediated temperature effects

In contrast to the effects of light, observed effects of temperature were primarily mediated through interactions with phytoplankton. Temperature had significant direct effects on bacterial community composition for SSB bacteria, but only marginally significant direct effects for TB bacteria. Lack of significant direct effects of temperatures spanning approximately 15°C on TB bacteria was surprising and may have been due to resource limitation. Alternatively, differences in the chemistry of the lake water added along with the bacterial treatment or the distribution of bacterial traits (e.g., ability to use light energy or breakdown available dissolved organic matter) may explain observed lake-specific differences in temperature response. Concentrations of dissolved organic carbon, total phosphorus, and total nitrogen tend to be higher in TB compared to SSB (Paver et al. 2013). Temperature effects were consistently significant in treatments with added phytoplankton, and enhanced in “away” phytoplankton treatments, potentially due to production of organic matter novel to the bacterial community (Fogg 1983, Sarmento and Gasol 2012). Bacterial community composition is continuously shaped by interactions with phytoplankton from their “home” lake (Paver et al. 2013), so initial bacterial assemblages were acclimated to “home” phytoplankton resources. Our results provide further evidence that bacteria rely on phytoplankton-derived dissolved organic carbon in darkly stained humic lakes with high background concentrations of dissolved organic carbon (Kritzberg et al. 2006, Kent et al. 2007).

Observed phytoplankton-dependent temperature effects along with previously published findings support a signal transduction hypothesis where changes in the environment are relayed to bacterial populations through their interactions with phytoplankton. In a suite of north temperate humic lakes (including SSB and TB), changes in the composition of phytoplankton assemblages were largely explained by environmental (e.g., water temperature, nutrients) and meteorological (e.g., photosynthetically active radiation, precipitation) factors (Kent et al. 2007). In contrast, changes in bacterial community composition were primarily explained by changes in phytoplankton population abundances and covariation between phytoplankton populations and the environment (Kent et al. 2007). Results from the current study provide experimental evidence that the signal of increased temperature in these lakes is largely relayed to bacteria by their interactions with phytoplankton. Experimental observations of Baltic Sea bacterial communities yielded a complementary result that phytoplankton bloom stage was a more important factor structuring bacterial communities than a 6°C change in temperature (Scheibner et al. 2013). Previous work on bacterial growth and activity provide additional support for environmental signal transduction via phytoplankton. Multiple regression and hierarchical partitioning analysis of data from 300 field studies indicated that temperature has a positive relationship with phytoplankton primary production, but not bacterial production (Faithfull et al. 2011). Despite positive correlations between temperature and bacterial production, bacterial production was primarily explained by a combination of total phosphorus and primary productivity (Faithfull et al. 2011). At low temperatures, bacterial growth and activity are frequently temperature limited (Simon and Wünsch 1998, Vrede 2005, Adams et al. 2010). When temperature is not limiting, bacterial growth in temperate lakes is commonly limited by phosphorus, dissolved organic carbon, or the two in combination (Vrede 2005). Phytoplankton clearly support bacterial growth and activity and have the potential to relay environmental signals to bacteria with which they interact.

### Temperature- and light- dependent of phytoplankton effects

Effects of phytoplankton on bacterioplankton were temperature, and to a lesser extent, light dependent. Observed interdependence of temperature and phytoplankton effects is consistent with the framework that phytoplankton provide organic matter resources to bacteria (Cole 1982, Sarmento and Gasol 2012) and temperature regulates the flow of carbon from phytoplankton to bacteria (Overmann 2013, Scheibner et al. 2013). Notably, when bacteria from TB were incubated with “home” phytoplankton, there was a significant interaction between light and temperature (Fig. 3). In contrast to high light assemblages that became increasingly different from their initial composition along a somewhat linear trajectory in ordination space as temperature increased, low light assemblages exhibited a curved response (Fig. 2). Bacterial composition in the coldest low light treatment was uncharacteristically different from the initial assemblage relative to other assemblages incubated at that temperature. These observations may be explained by changes in the concentration and composition of phytoplankton exudates under different temperature and light conditions. Combined effects of light and temperature have been shown to affect the dominant metabolic pathways used to process carbon, thereby controlling exudate release (Parker and Armbrust 2005). Alternatively, the dominant mechanism of bacteria-phytoplankton interactions may change depending on light and temperature. For example, low light conditions can increase the rate of bacterial consumption by mixotrophic phytoplankton under certain conditions (e.g., low nutrient availability; [McKie-Krisberg and Sanders 2014]).

### Consequences for planktonic microbial communities

Observations that light has primarily direct effects on bacterioplankton, while temperature effects are mediated through interactions with phytoplankton, have implications for interpreting seasonal changes in planktonic communities and forecasting future changes. The pronounced, direct effect of light on bacterial community composition emphasizes the importance of collecting high-resolution samples over depth and incorporating mechanisms structuring bacterial communities into a depth-specific framework. Temperature is frequently correlated with the succession of aquatic bacterial communities (Crump and Hobbie 2005, Fuhrman et al. 2006, Shade et al. 2007). Our results demonstrate that, in some systems, much of the bacterial community response to temperature is fueled by their interactions with phytoplankton. Thus, if the objective is to forecast how bacterial community structure and function will change in response to environmental changes, it is necessary to incorporate the predicted response of phytoplankton and context-dependence of interactions linking phytoplankton and bacterial assemblages. For example, elevated temperature in mesocosms during the spring phytoplankton bloom in Kiel Bight accelerated the onset of the phytoplankton bloom, decreased the intensity of maximum chlorophyll a and particulate organic carbon by approximately 20%, and caused dissolved organic carbon concentrations to increase more rapidly than under ambient temperature conditions (Biermann et al. 2014).

Environmental context is critical for understanding how planktonic communities will change over time and in response to environmental change. Many of our observations were highly dependent on the biotic or environmental context – responses seen in one lake were not replicated in the other and, for TB bacteria combined with TB phytoplankton, response to temperature depended on light availability. Context-dependence has also been described for bacterial production in high mountain lakes where bacterial response to solar radiation treatments depended on the presence of phytoplankton and whether bacteria were phosphorus-limited (Medina Sánchez et al. 2002). The prevalence of non-additive interaction effects among environmental factors emphasizes the importance of multi-factorial experimental investigations into drivers of microbial community composition and activity. It additionally underscores a need to identify mechanisms underlying context-dependent microbial responses and build these mechanisms into frameworks describing aquatic microbial community dynamics.

### Effects of interacting factors on community composition

Phytoplankton, light availability, and temperature act in concert to bring about changes in bacterial community composition over time. Light availability directly affects bacterial community composition while interactions with phytoplankton amplify or relay the signal of increasing temperature to bacteria. The strength of phytoplankton interactions with bacteria, inferred through comparisons of bacterial community composition across temperature and light treatments relative to no-phytoplankton controls, depends on temperature. For certain combinations of phytoplankton and bacteria, the outcome of phytoplankton interactions appears to additionally be light dependent. The enhanced effect of “away” phytoplankton relative to “home” phytoplankton and the lack of consistent temperature effects in treatments without phytoplankton provide strong support for phytoplankton resources shaping bacterial communities and their response to environmental conditions. These findings emphasize the importance of observing population and community responses to multiple ecological drivers simultaneously and under a range of environmentally relevant conditions. Studies aimed at understanding the effects of climate change frequently compare ambient temperature conditions to an elevated temperature treatment. Our results suggest that there are inflection points in community responses to temperature that would be overlooked in ambient vs. elevated temperature comparisons. The idea that under-sampling the range of potential temperatures constrains our ability to make general inferences about temperature effects is reinforced by studies that have investigated the effect of temperature on bacterial production and observed two temperature optima (Simon and Wünsch 1998, Adams et al. 2010). The problem of under-sampling treatment levels is potentially problematic for other factors as well, including light availability (Gu and Wyatt 2016). Movement away from context-specific observations towards generalizable, theoretical advances and identifying parameters that can be incorporated into predictive frameworks will depend on investigations such as this into the mechanisms driving observed changes in community composition.

## Acknowledgements

We thank E. Baird, B. Crary and K. Hayek for assistance carrying out the experiment; K. Hayek for lab assistance; K. McMahon lab and the staff of UW-Madison Trout Lake Research Station for logistical support; C. Cáceres, W. Metcalf, A. Peralta, R. Whitaker and A. Yannarell for constructive feedback on earlier versions of this manuscript. Funding was provided by NSF grant MCB-0702653 to A.D.K. and NSF DDIG grant DEB-11-10623 DISS to S.F.P.

